# Quantitative Head-to-Tail Analysis Reveals Heterogeneous Cellular Organization and Connexin Expression in the Porcine Sinoatrial Node

**DOI:** 10.64898/2026.11.05.764659

**Authors:** Hillary K. J. Kao, Regan L. Smithers, David H. Ramos-Rodriguez, Kaitlin Van Brusselen, Rableen K. Kaleka, David Cameron, Jan A. Nolta, Kyle Fink, J. Kent Leach, L. Fernando Santana, Nipavan Chiamvimonvat, Deborah K. Lieu

**Affiliations:** Department of Internal Medicine, Division of Cardiovascular Medicine, University of California, Davis, Sacramento, CA, 95817, USA; Institute for Regenerative Cures and Stem Cell Program, University of California, Davis, Sacramento, CA 95817, USA; Department of Biomedical Engineering and Orthopedic Surgery, University of California, Davis, Sacramento, CA, 95817, USA; Bridges to Stem Cell Research Program, California State University, Sacramento, Sacramento, CA, 95819, USA; Department of Neurology, University of California, Davis, Sacramento, CA, 95817, USA; Department of Physiology and Membrane Biology, University of California, Davis, Davis, CA 95616, USA; Department of Basic Medical Sciences, Department of Internal Medicine, and Translational Cardiovascular Research Center, University of Arizona College of Medicine - Phoenix, Phoenix, AZ, 85004, USA

## Abstract

**Background:** The sinoatrial node (SAN) is an integral component of the cardiac conduction system that dictates heart rhythm. Optical mapping studies have demonstrated functional differences between the superior and inferior leading pacemaking sites in the human SAN; however, the mechanistic connections between structure and function in the SAN through intra-tissue and inter-tissue current source-sink relationships have not been fully resolved.

**Methods:** Porcine SAN longitudinal cryosections along the head-to-tail axis were immunostained and analyzed for regional cell organization and connexin (CX) isoform expression.

**Results:** Quantification of pacemaking cardiomyocyte (PCM) density revealed >9-fold more PCMs within a SAN volume than suggested by a widely cited estimate, assuming comparable PCM densities across species. The SAN head region consisted of large PCM clusters with low aspect ratios, high PCM density, low fibroblast density, and PCMs with the highest HCN4 expression and lowest CX40 and CX43 expression. These characteristics are consistent with an electrically well-insulated, strong current-source that supports pacemaking function. Although the SAN tail also exhibited high local PCM density, its smaller PCM clusters, lower HCN4 expression, and higher CX40, CX43, and CX45 expression rendered it a weaker current source. The SAN mid-region exhibited large PCM clusters aligned in the head-to-tail axis and the highest expression of CX40 and CX45, an expression profile that suggests a fast-conducting intra-tissue pathway that electrically links the leading pacemaking sites at the head and tail. The peripheral SAN at the head and tail also exhibited higher CX expression of all isoforms than the central subregions, an organization consistent with enhanced inter-tissue coupling required for proper pacing function.

**Conclusions:** Our data provide novel insights into the distinct regional micro-organization of the SAN along the head-to-tail axis, which through intra-tissue and inter-tissue coupling, likely modulates the balance of current source-sink relationships that supports robust pacemaking and pacing functions.

## INTRODUCTION

The sinoatrial node (SAN) is responsible for regulating cardiac sinus rhythm. Shaped like a comma, the SAN lies along the crista terminalis (CT), with its head near the superior vena cava (VC) and its tail near the inferior VC.^1^ This pacemaking tissue is structurally and functionally distinct from the surrounding contractile tissue composed of atrial cardiomyocytes (ACMs). The SAN consists of predominantly pacemaking cardiomyocytes (PCMs) and fibroblasts (FBs) embedded within a highly fibrous extracellular matrix. PCMs function as the current source that sets the heart rate by initiating spontaneous action potentials. These parenchymal cells are well-supported by surrounding FBs that maintain the pacemaking microenvironment through production of growth factors and matrix proteins; however, because FBs are non-excitable, they can also act as a current sink and suppress pacemaking function.^2^ The organization of these major SAN components dictates intra-tissue (PCMs to PCMs and FBs) and inter-tissue (PCMs to ACMs) electrical couplings, establishing an optimal source-sink relationship that is essential for proper pacemaking and pacing functions. Details of how this organization maintains an optimal current source-sink relationship remain to be elucidated.

Optical mapping of rat and human SANs have revealed the presence of two leading pacemaking sites, one located at the superior SAN and another at the inferior region.^3,4^ Notably, the superior pacemaking site at the SAN head exhibits higher frequency of automaticity (>81 bpm) than the inferior site (<66 bpm), suggesting regional differences in cellular phenotype and micro-organization that warrant further investigation. Regional organization along this axis has precedent in other species, where microvascular density and myocyte firing rate differ between the superior and inferior node.^5^ To date, histology of the SAN has been conducted almost exclusively in transverse-oriented tissue sections, limiting spatial correlation with optical mapping along the head-to-tail axis.^4,6–10^ Additionally, histological staining, such as hematoxylin & eosin or Masson’s trichrome, provides only high-level anatomy with little information on the spatial arrangement of current source-sink components.

Our previous study qualitatively revealed distinct regional intra- and inter-tissue micro-organization, as well as differential regional expression of the pacemaking hyperpolarization-activated cyclic nucleotide-gated channel (HCN4) and T-Box Transcription Factor 18 (TBX18) genes in PCMs along the head-to-tail axis in large mammalian SANs.^11^ The structural and molecular differences across SAN regions improved our understanding of why the leading pacemaking sites localize predominantly to the superior and inferior SAN; however, many questions remain, including the number of resident PCMs and FBs, their densities and spatial organization, and the cellular gap junction expression that affects the source-sink relationship in the SAN. Building on our prior work, the present study quantitively characterizes the regional PCM cluster architecture and cellular composition, as well as gap junction expression in PCMs and FBs along the head-to-tail axis of the porcine SAN. Our findings highlight plausible contribution of the distinct regional SAN structure and electrical coupling to pacemaking and pacing functions, providing foundational insights for SAN physiology and tissue organizational criteria for engineering biopacemakers.

## METHODS

### Immunostaining and Imaging of Porcine Sinoatrial Nodes

Fresh 6-month-old market porcine hearts were acquired from the UC Davis Meat Laboratory. SAN tissues were manually dissected and embedded in Tissue-Tek O.C.T compound. Tissues were sectioned at 30-μm thickness using a Thermo Fisher Cryotome and mounted onto Superfrost plus slides. Longitudinal cryosections ∼300-1000 μm from the epicardial surface were immunostained after fixation with 4% (w/v) paraformaldehyde and permeabilized with 0.3% (v/v) Triton X-100. To determine pacemaking cluster dimensions and cell density, cryosections from three pig hearts were immunostained with cardiac troponin T (TNT) to identify CMs, HCN4 to identify PCMs, and vimentin (VIM) for FBs. Connexin (CX) expression was evaluated in the cells by co-staining CX40, CX43, or CX45 with HCN4 and VIM. All cryosections were nuclear counterstained with DAPI. Primary and secondary antibody dilution factors are described in **Table S1**. All images were captured with either a Zeiss LSM 900 Airyscan2 confocal microscope or Leica STELLARIS confocal microscope from three pig hearts and analyzed using Imaris software version 10.1 (Oxford Instruments) as described below.

### Regional PCM Cluster and Cell Organization Analysis

The SAN was identified as the HCN4^+^/TNT^+^ region and manually delineated from the neighboring VC and CT. PCM structures were generated by identifying HCN4^+^ clusters using the trainable machine learning segmentation tool in Imaris and subsequently assessed for individual cluster properties. Clusters with an area less than 320 μm^2^, the approximate area for one PCM^1^, were filtered out to exclude partial cells in the PCM cluster analysis. Total HCN4 area (%) was calculated as the summed HCN4^+^ area divided by the total area in each SAN region. Aspect ratio of the PCM clusters was calculated as the ratio of length to width of the defined object-oriented bounding box surrounding the HCN4^+^ cluster area.

PCM and FB densities were quantified by counting the number of nuclei within the PCM (HCN4^+^) and FB (HCN4^−^) areas, respectively. Nuclei were rendered as spots using a 6-μm diameter filter and an intensity quality threshold of 8. Local PCM density was calculated as the total number of PCMs within the total HCN4^+^ area. Local FB density was calculated as the total number of FBs within the total HCN4^−^ area. Regional PCM and FB densities within the SAN head, middle, and tail regions were quantified as the total number of respective PCMs and FBs within the total regional area (HCN4^+^ and HCN4^−^ tissue combined).

### Regional CX Expression Analysis in PCMs and FBs

To quantify CX expression of isoforms CX40, CX43, and CX45, image fields of 240 µm x 240 µm from three hearts were manually selected from the central and peripheral subregions within each of the SAN head, middle, and tail regions, yielding six SAN subregions for analysis per porcine heart. Corresponding image fields from the VC and CT served as atrial controls. SAN PCM and ACM structures, generated as described above, were used to create masks for quantifying CX expression in the PCM clusters. CX fluorescence detected within HCN4-high and HCN4-low structures was regarded as expression levels in PCMs or ACMs, respectively. Structures were generated for CX puncta to quantify mean intensity values and puncta plaque area. CX puncta mean fluorescence intensity and area data were log_2_-transformed and presented as frequency histograms. To analyze CX expression in FBs, FB surfaces were generated by identifying VIM^+^ cells using VIM fluorescence intensity. Partial FBs with a cell body area <20 μm^2^ or no detected nucleus were excluded from analyses. CX expression in FBs was quantified by fluorescence intensity within VIM^+^ structures due to limitation in detecting individual puncta within the small cell size. Mean fluorescence intensity for all cell types in each region was normalized to corresponding FB mean fluorescence intensity in the CT region.

### Statistical Analysis

Results are presented as mean ± SEM or median ± IQR as specified. Statistical analyses were performed using GraphPad Prism (version 11). Data was assessed for normality using a D’Agostino and Pearson Test or Shapiro-Wilk Test with a = 0.05. Statistical significance was determined by one-way ANOVA with Tukey’s post hoc test or Kruskal-Wallis using Dunn’s post hoc test as appropriate. P-values are denoted as \**P* < 0.05, \*\**P* < 0.01, \*\*\**P* < 0.001, \*\*\*\**P* < 0.0001.

## RESULTS

### Regional Cellular Organization in the SAN

We previously reported, qualitatively, distinct regional micro-organization along the head-to-tail axis of the SAN.^11^ Here, we quantitatively characterized the organizational properties of these regions using detailed image analysis. PCMs were identified as TNT^+^/HCN4^+^ cells and FBs as VIM^+^ cells (Figure 1A, Figure S1). The HCN4^+^ SAN, traversed by the SAN artery, was flanked laterally by the CT and VC, arching toward the VC at the superior end. To enhance visualization of regional differences along the head-to-tail axis, HCN4 expression was presented in pseudo-color based on fluorescence intensity, revealing highest HCN4 expression in the SAN head close to the superior VC, followed by high HCN4 expression in the SAN tail close to the inferior VC, and low expression in the SAN mid-region (Figure 1B, Figures S4-S6). Unlike the abrupt transition in HCN4 expression at the SAN-VC interface, the SAN-CT interface exhibited a HCN4 expression gradient, decreasing towards the CT. The SAN was segmented into three regions by HCN4 expression and PCM cluster organization. PCM clusters were delineated as structures based on HCN4 fluorescence intensity, then each PCM cluster was rendered in pseudo-color based on the mean HCN4 intensity (Figure 1C). PCM clusters in the SAN exhibited distinct HCN4 expression— with the highest cluster mean HCN4 fluorescence in the SAN head, followed by the SAN tail and then the mid-region—allowing clear division of the SAN into head, middle, and tail regions. Notably, the three regions consisted of distinct cluster organization. The SAN head and tail displayed minimal preferential cluster arrangement, in stark contrast to the aligned PCM clusters in the mid-region spanning the two ends (Figure 1D).

**Figure 1.**
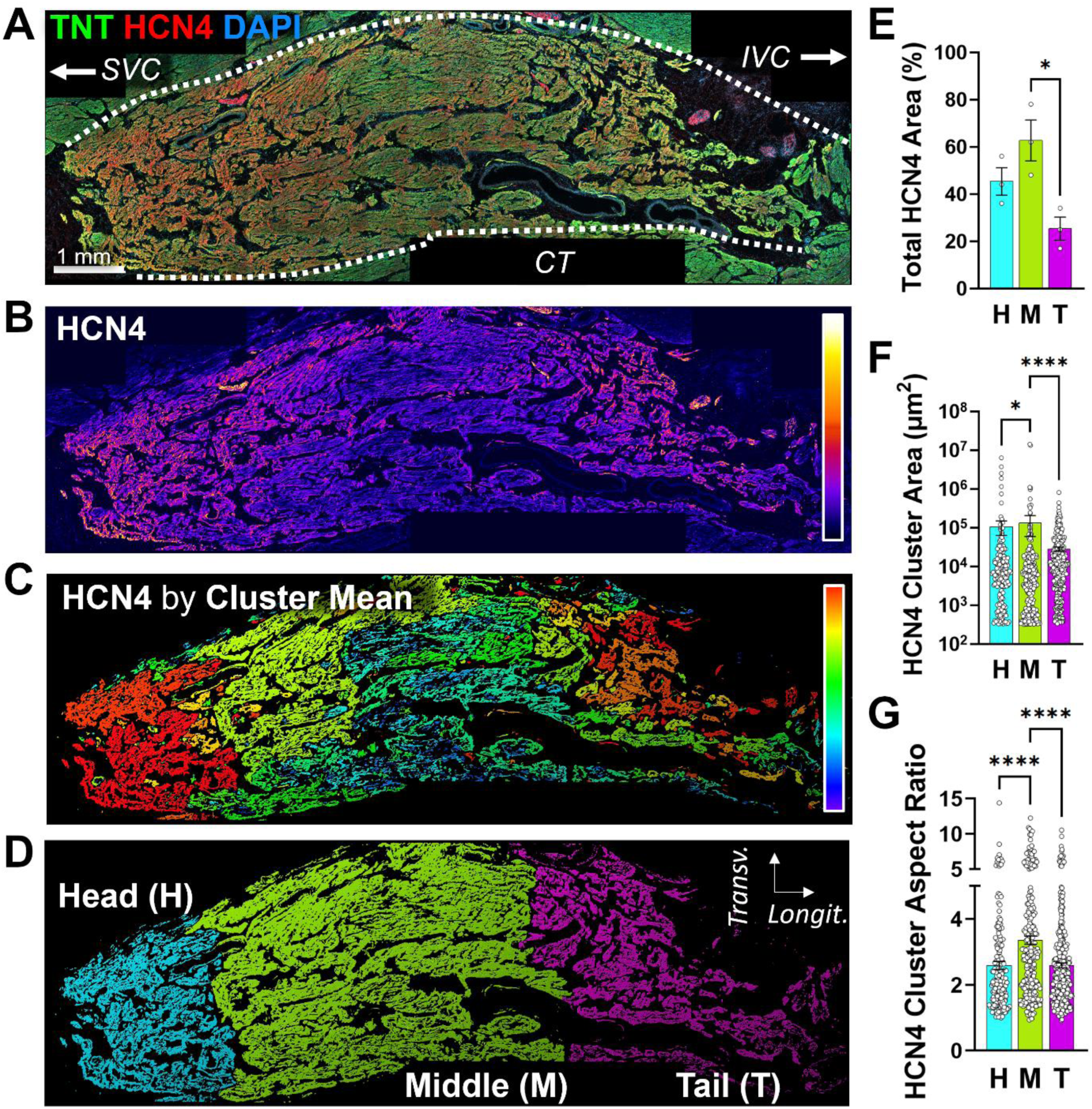
PCM cluster organization in the porcine SAN. **A)** Representative SAN cryosection immunostained for HCN4 and TNT, showing cellular organization from the head to tail. **B)** Pseudocolor image of the SAN based on HCN4 mean intensity. **C)** Pseudocolor image of the SAN based on mean HCN4 intensity within each PCM cluster. **D)** Pseudocolor image of the SAN segmented longitudinally into head (H), middle (M), and tail (T) regions based on HCN4 expression and cluster organization. **E)** Quantification of total HCN4^+^ area in each SAN region. **F)** Quantification of HCN4^+^ cluster area within individual PCM clusters in each SAN region. **G)** Quantification of the aspect ratio for individual HCN4^+^-PCM clusters. All quantifications were derived from three hearts, one image from each heart. \**P* < 0.05, **** for *P* < 0.0001. (TNT: troponin T, HCN4: hyperpolarization-activated cyclic nucleotide-modulated channel 4, SVC: superior vena cava, IVC: inferior vena cava, CT: crista terminalis, Transv: transverse, Longit: longitudinal, H: head region, M: middle region, T: tail region)

PCM coverage was quantified as the fraction of the SAN region occupied by the HCN4^+^ structures. Quantitative analysis revealed PCMs in the mid-region occupied a large fraction of the SAN, comprising approximately ∼63% of the total tissue area, compared to ∼45% in the head and ∼25% in the tail region (Figure 1E). The area of individual PCM clusters was the largest in the mid-region at 1.4 x 10^5^ µm^2^, which was ∼1.2-fold and ∼4.7-fold higher than the head and tail, respectively (Figure 1F). The aspect ratio of PCM clusters within the mid-region was the highest at ∼3.3, which was ∼1.3-fold more elongated than both the head and tail regions (Figure 1G). Collectively, quantifications of porcine SANs demonstrated distinct spatial patterns in PCM architecture, characterized by large, less aligned PCM clusters in the head, small and equally less aligned clusters in the tail, and largest, highly directional, and elongated PCM clusters in the middle region that bridged the two ends.

### Regional Cellular Density in the SAN

To assess the cellular composition of the SAN, PCM and FB densities in the head, middle, and tail regions were quantified in tissue sections stained for HCN4, VIM and DAPI (Figure 2A, 2B). PCM and FB counts were quantified by the number of nuclei within the respective HCN4^+^ and HCN4^−^ areas (Figure 2C). The total cell density, combining PCMs and FBs in the SAN for all regions, exhibited comparable mean cell density, ranging from ∼131,000 to 134,000 cells/cm^2^ (Figure S3). Of note, SAN sections were stained for CD31, HCN4, VIM, and DAPI to assess the fraction of CD31^+^-endothelial cells (ECs) that were embedded in the HCN4^+^ or VIM^+^ structures (Figure S2). Misassignment of ECs as PCMs and FBs accounts for ∼5% of cells within the HCN4^+^-PCM area and ∼10% of the cells within the VIM^+^-FB area, respectively. Notably, SAN microvascular density in mice is graded along the head-to-tail axis.^5^ Nevertheless, ECs are expected to inflate the SAN cell density only modestly by ∼5–10%, although the magnitude of this inflation may vary along the axis.

**Figure 2.**
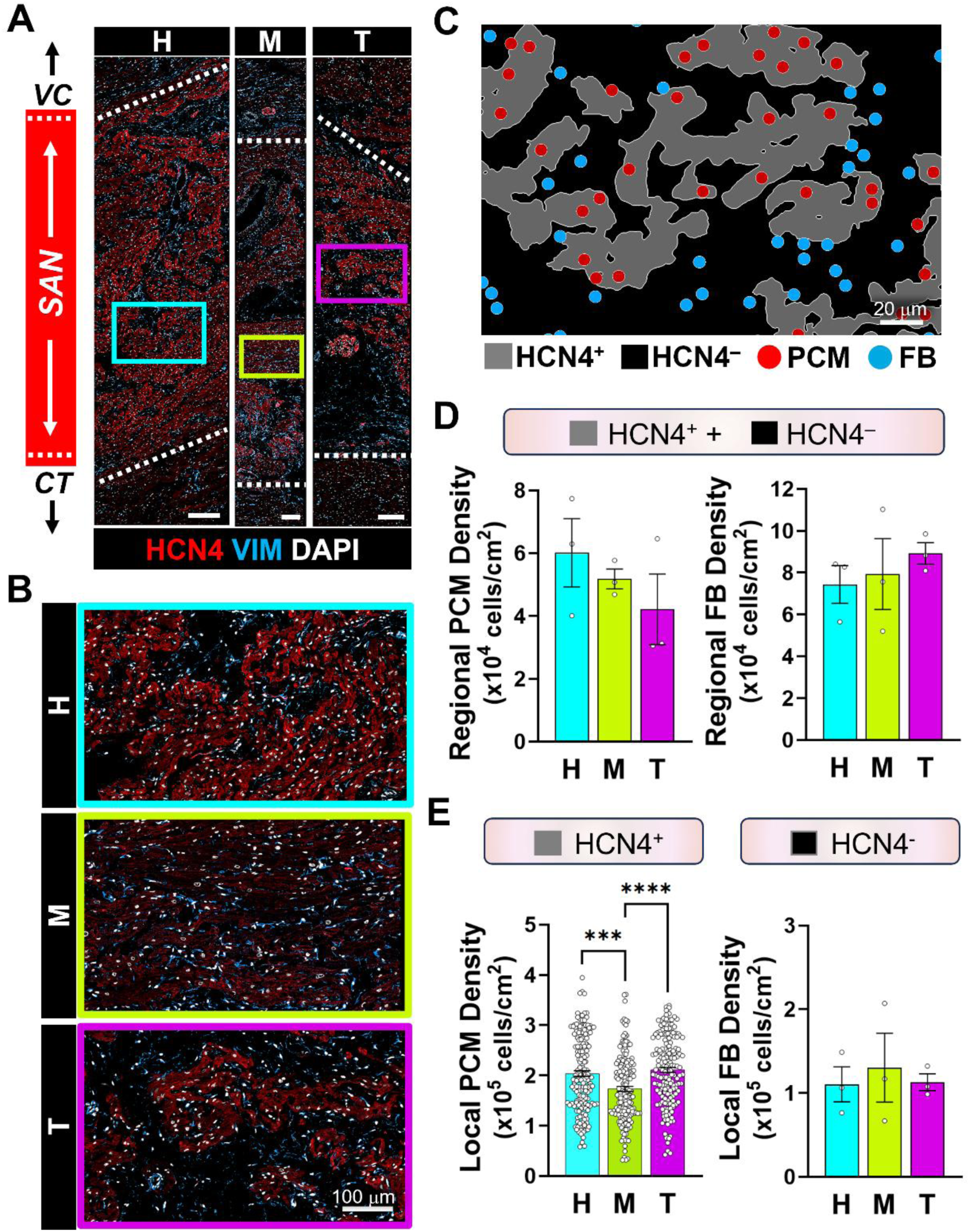
PCM and FB cell density in the porcine SAN. **A)** Images of the SAN head (H), middle (M), and tail (T) regions immunostained with HCN4 and VIM to identify PCMs and FBs, respectively (scale bar = 200 µm). **B)** Magnified images of cellular organization for regions boxed in panel A. **C)** Imaris rendering of the SAN showing HCN4^+^ masks delineating PCM clusters and spots marking PCM and FB nuclei. **D)** Quantified regional PCM and FB densities in each SAN region. **E)** Quantified local PCM and FB densities within HCN4^+^ and HCN4^-^ area, respectively, in each SAN region. All quantifications were derived from three images, one from each heart. *** *P* < 0.001, **** *P* < 0.0001.

Regional PCM and FB densities were quantified by the nuclear count within combined HCN4^+^ and HCN4^−^ areas in each of the SAN regions (Figure 2D). The regional PCM density in the SAN head was ∼60,000 cells/cm^2^, higher than the middle and tail regions by ∼14% and 30%, respectively, albeit without statistical significance. Conversely, regional FB density followed the opposite trend, with the highest density of ∼89,000 cells/cm^2^ in the tail region that was higher by ∼17% in the head and 11% in the middle region, without statistical significance. Local PCM and FB densities were quantified by the nuclear count within the respective HCN4^+^ or HCN4^−^ area in each of the SAN regions, revealing a PCM density of ∼204,000 cells/cm^2^ in both the head and tail regions that was statistically higher than the mid-region at ∼165,000 cells/cm^2^ (Figure 2E). The local FB density outside PCM clusters (HCN4^−^ area) in the SAN head was ∼110,000 cells/cm^2^, trending lower by 15% and 3% than middle and tail regions, respectively. Interestingly, both regional and local PCM densities in the SAN regions exhibited inverse trends to those of regional and local FB densities, such that regions of high PCM density coincided with low FB density.

### Differential CX Expression along the SAN Head-to-Tail Axis

To examine gap junction distribution along the head-to-tail axis in the SAN, cryosections were immunostained for CX40, CX43, or CX45 in conjunction with HCN4 and VIM to identify PCMs and FBs, respectively. Since HCN4 and CX expression differed between the central and peripheral SAN, the head, middle, and tail regions were further subdivided to yield 6 subregions. CX expression in PCM and ACM was quantified for puncta density, puncta fluorescence intensity, and puncta plaque area, with histogram distribution shown for the latter two parameters. Puncta with intensity or area greater than three standard deviations above the mean across all cardiac regions were identified as rare large plaques formed by numerous aggregating gap junction proteins. CX expression in FBs was measured as total mean CX fluorescence due to unresolvable individual puncta at the image resolution.

CX40 was present in both FBs and PCMs in the SAN (Figures 3A, 3B; Figures S4). In FBs, CX40 fluorescence was lowest in the SAN mid-region, at ∼17% lower than the SAN head and ∼12% lower than the SAN tail and atrial regions (Figure 3C). Comparing CM subtypes, PCMs and ACMs generally had comparable CX40 puncta density with ∼57-60 puncta/1000 μm², except for PCMs in the SAN mid-central and tail-peripheral subregions with higher puncta density respectively at ∼77 and ∼80 puncta/1000 μm² (Figure 3D), suggesting high electrical coupling between PCMs in the mid-central subregion connecting the SAN head and tail, as well as improved conductance between PCMs at the tail-periphery forming conduction exit paths. CX40 fluorescence intensity histogram distributions were right-skewed with the same intensity peak in PCMs of all subregions; however, puncta intensity distribution was the broadest in ACMs followed by PCMs in the SAN mid-central subregion, which was attributed to CX40 clusters forming large plaques with high fluorescence intensity (Figure 3E). Indeed, these rare plaques accounted for ∼11% of all puncta in ACMs and ∼7% of puncta in mid-central PCMs, contrasting with merely ∼0.5-3% large plaques in PCMs of other subregions. CX40 puncta area distributions, peaking near ∼0.3 μm², followed the same intensity distribution pattern for all subregions, which is consistent with correlation between puncta intensity and size. As expected, ACMs contained the greatest proportion of large plaques ≥ 2 μm² at ∼4%, followed by ∼2% of puncta in the mid-central PCMs and ≤ 1% puncta in PCMs of all other subregions (Figure 3F). The maximum puncta area in the atrial region was ∼40 μm², in contrast to the central mid-subregion at ∼16 μm², and the peripheral head and tail subregions at ∼12 μm² and < 8 μm² for the remaining subregions. Interestingly, mid-central PCMs were aligned and expressed high intensity, large CX40 aggregates that colocalized with HCN4 at the intercalated discs similar to ACMs (Figure S7), suggesting contribution of CX40 gap junctions to anisotropic conduction directed in the head-to-tail axis. Considering the expression, preferential location, and high single-channel conductance of CX40 in the SAN, central-middle and peripheral-tail subregions identified as the CX40-rich support intra-tissue electrical coupling for rapid conduction between the head and tail and inter-tissue coupling for potential exit pathways to the atrial myocardium.

**Figure 3.**
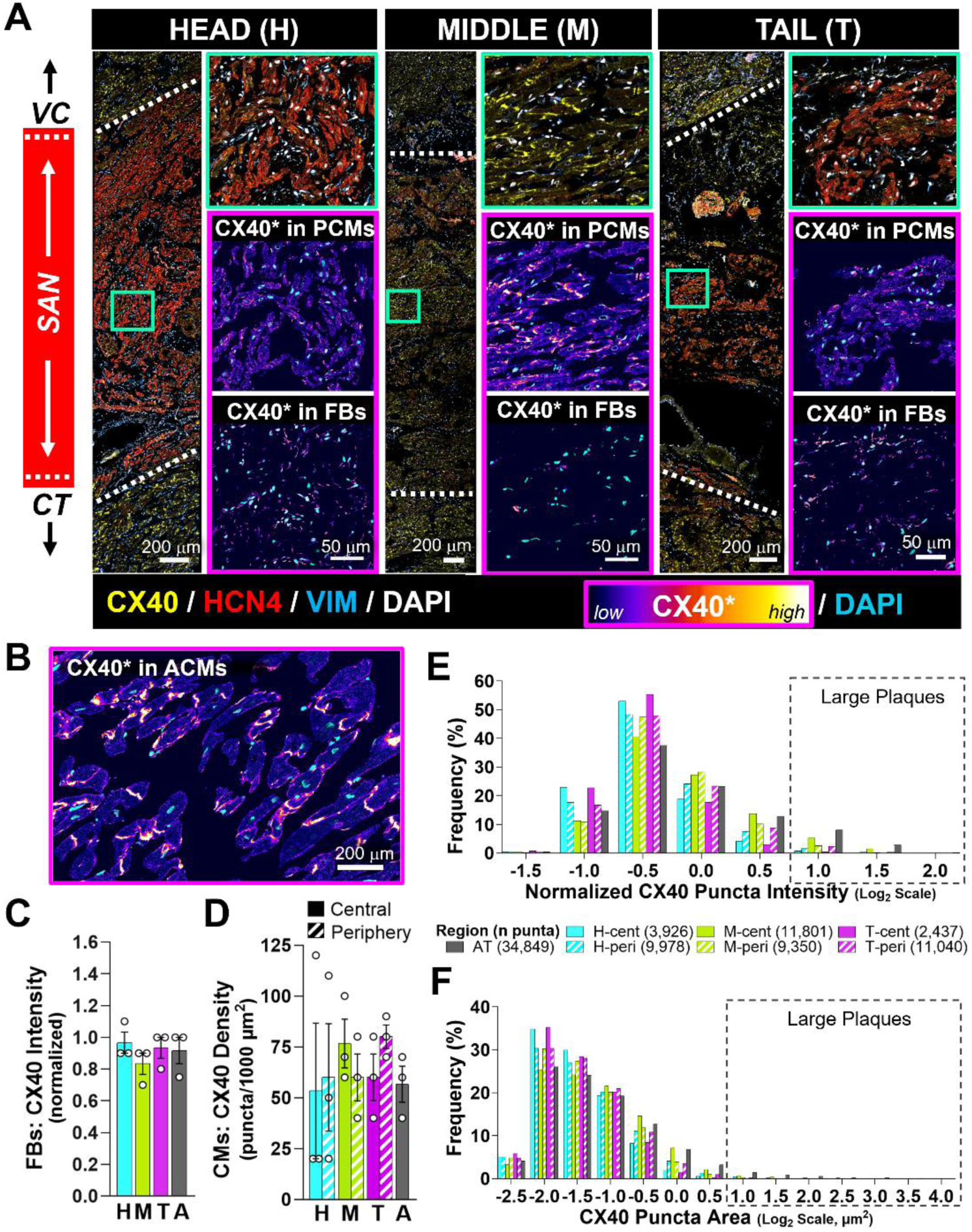
Distribution of CX40 in the porcine SAN. **A)** Representative immunofluorescence images of the SAN head, middle, and tail regions immunostained for CX40, HCN4, VIM, and DAPI. White dashed lines indicate the SAN-atrial border, based on HCN4 expression. Insets show a magnified central subregion with corresponding pseudocolor images for CX40 expression in PCMs and FBs. **B)** Representative pseudocolor image for CX40 expression in ACMs. **C)** Normalized CX40 expression in FBs within the SAN and atrial (A) regions. **D)** CX40 puncta density in CMs within SAN and A regions. **E)** Distribution of CX40 puncta fluorescence intensity for 6 SAN subregions and atrial tissue (AT). **F)** Distribution of CX40 puncta area for the same subregions. Outlined boxes highlight CX40 plaques with high fluorescence intensity and large area. All quantifications were pooled from three hearts, one 240 μm x 240 μm image from each heart. (H: head, M: middle, T: tail, A: atrial, central: peri: peripheral, AT: atrial tissue; cent)

CX43, as expected, was detected in ACMs, and notably also throughout the SAN (Figures 4A, 4B; Figures S5, S8). FBs in the neighboring atrium expressed lower CX43 compared to those in the SAN, with ∼10-20% higher expression in the tail-FBs than the other subregions, although without statistical significance (Figure 4C). Comparing CMs across the SAN and bordering atrial tissue, CX43 puncta density was highest in the atrium with ∼87 puncta/1000 μm² and SAN tail-peripheral subregion with ∼93 puncta/1000 μm² but much lower in the head-central subregion at ∼27 puncta/1000 μm² (Figure 4D). Notably, CX43 puncta were also consistently denser in the peripheral than central SAN subregions, especially at the SAN head and tail, which may mark conduction exit pathways. CMs in all tissues exhibited right-skewed distributions of CX43 puncta intensity and area, however this trend was most pronounced in ACMs. Large CX43 puncta plaques ≥ 1.4 μm² constituted ∼8% of the total puncta in the atrial region with puncta area up to ∼59 μm², but only 0.3-1.5% of the puncta in all SAN subregions with a maximum puncta area of 2-13 μm². This high puncta density in the atrial region in conjunction with large CX plaques indicate substantially higher atrial conductance via CX43 gap junctions than that in the SAN (Figures 4E, 4F). Like CX40 in the atrial tissue, high-intensity and large CX43 plaques also exhibited colocalization with HCN4 at intercalated discs (Figure S8); however, those CX43 aggregates were largely absent in the mid-central SAN subregion (Figure 4A). This is reflected in the differing puncta distribution pattern in the SAN mid-region between CX43 and CX40, with only ∼1% rare large CX43 plaques. PCMs in the head- and tail-peripheral subregions also exhibited right-shifted puncta intensity and area distribution peaks, with ∼1-2% rare large and high-intensity CX43 plaques, possibly providing electrical coupling in conduction exit pathways at the atrial-SAN interface in those regions.

**Figure 4.**
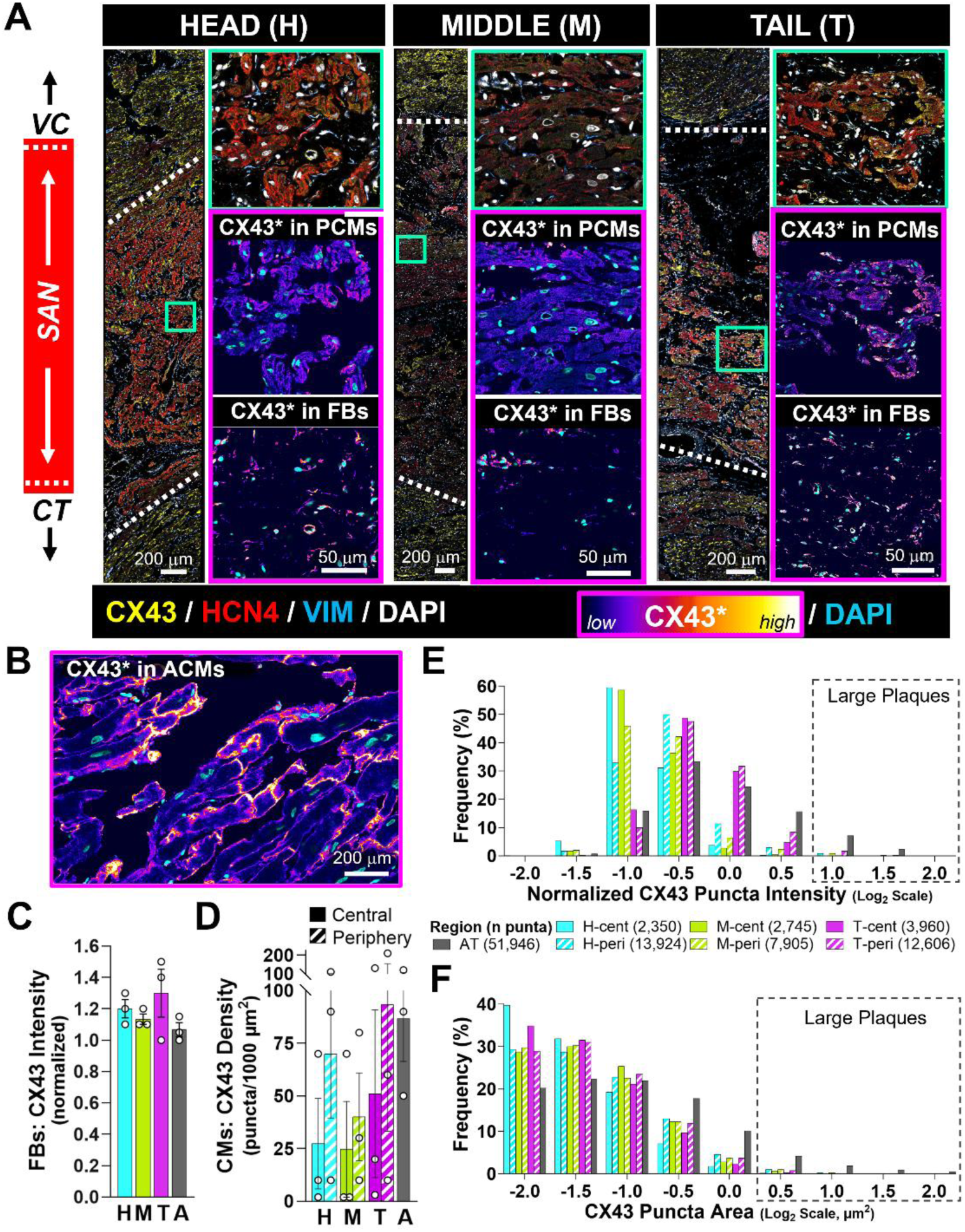
Distribution of CX43 in the porcine SAN. **A)** Representative immunofluorescence images of the SAN head, middle, and tail regions immunostained for CX43, HCN4, VIM, and DAPI. White dashed lines indicate the SAN-atrial border, based on HCN4 expression. Insets show a magnified central subregion with corresponding pseudocolor images for CX43 expression in PCMs and FBs. **B)** Representative pseudocolor image for CX43 expression in ACMs. **C)** Normalized CX43 expression in FBs within the SAN and atrial (A) regions. **D)** CX43 puncta density in CMs within SAN and A regions. **E)** Distribution of CX43 puncta fluorescence intensity for 6 SAN subregions and atrial tissue (AT). **F)** Distribution of CX43 puncta area for the same subregions. Outlined boxes highlight CX43 plaques with high fluorescence intensity and large area. All quantifications were pooled from three hearts, one 240 μm x 240 μm image from each heart. (H: head, M: middle, T: tail, A: atrial, central: peri: peripheral, AT: atrial tissue; cent)

Small CX45 puncta were dispersed throughout the SAN and atrial myocardium (Figures 5A, 5B; Figures S6, S9). CX45 expression was comparable in FBs across the head, middle, and atrial regions but ∼14% lower in the tail (Figure 5C). Unlike CX40 and CX43, CX45 puncta did not colocalize with HCN4 at the intercalated discs in atrial tissue (Figure S9), nor did they form large plaques in the intercalated discs in the mid-SAN region (Figure 5A). The presence of CX45 in ACMs, albeit at low density of ∼57 puncta/1000 μm², does not support exclusive CX45 localization to the SAN; however, CX45 puncta density was generally higher in PCMs than ACMs (Figure 5D). Of the PCMs in the SAN subregions, those in the SAN mid-central subregion had the highest CX45 puncta density at ∼173 puncta/1000 μm², whereas those in the head-central subregion expressed the lowest density at ∼83 puncta/1000 μm², further suggesting weak conductance in the central head subregion. Overall, the CX45 puncta density in PCMs was nearly double that of CX40 and CX43. In contrast with the higher single-channel conductance CX isoforms, CX45 puncta intensity and area histograms showed comparable right-skewness and distribution peaks across all subregions (Figures 5E, 5F). While CX45 puncta ranged in size from ∼0.2-6 μm², the majority (> 70%) measured ∼0.3-0.4 μm² in all subregions. Hence, the subregional CX45 expression differences were mainly attributed to the puncta density rather than rare large plaques that make up ≤ 0.2% of the total puncta in all subregions.

**Figure 5.**
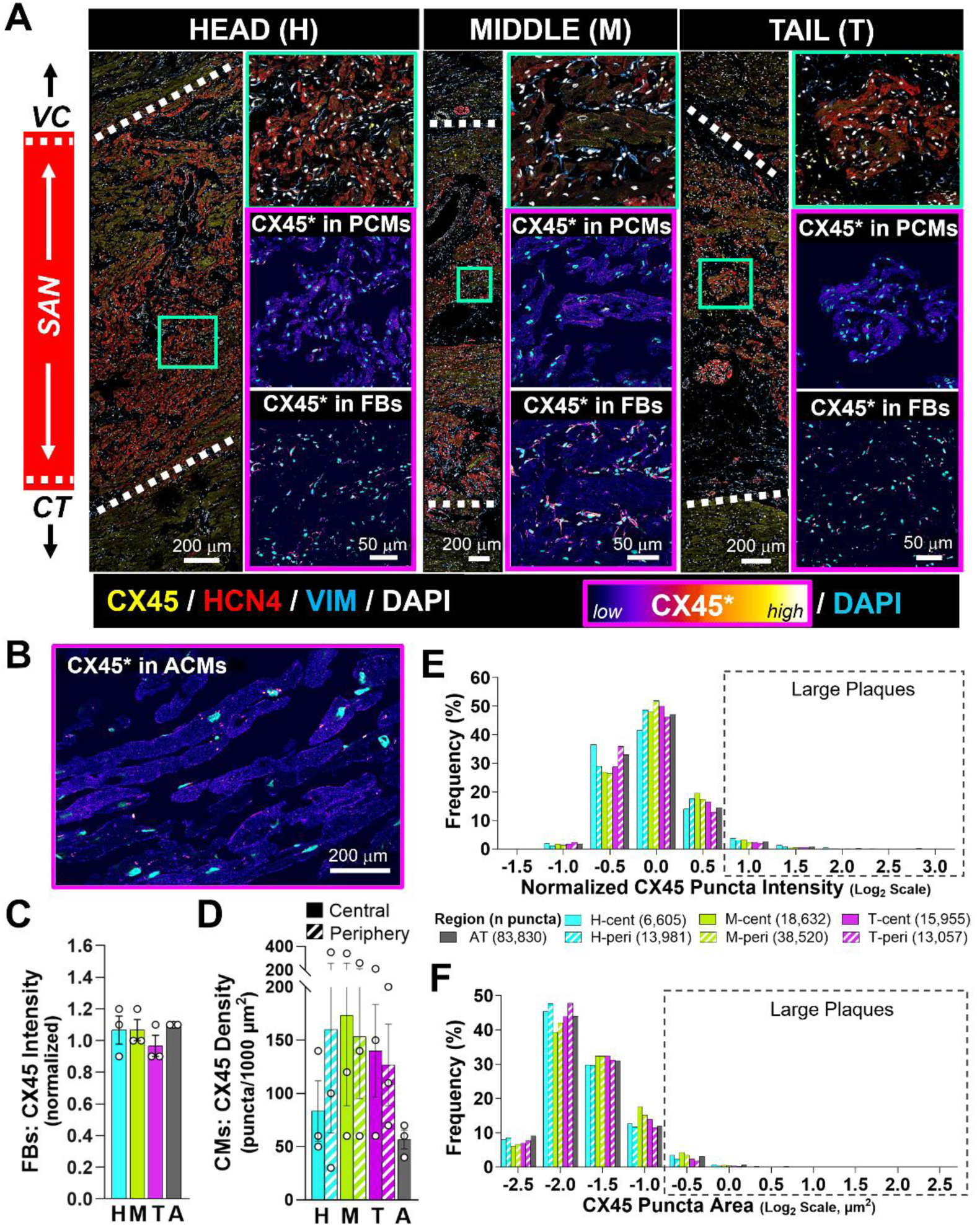
Distribution of CX45 in the porcine SAN. **A)** Representative immunofluorescence images of the SAN head, middle, and tail regions immunostained for CX45, HCN4, VIM, and DAPI. White dashed lines indicate the SAN-atrial border, based on HCN4 expression. Insets show a magnified central subregion with corresponding pseudocolor images for CX45 expression in PCMs and FBs. **B)** Representative pseudocolor image for CX45 expression in ACMs. **C)** Normalized CX45 expression in FBs within the SAN and atrial (A) regions. **D)** CX45 puncta density in CMs within SAN and A regions. **E)** Distribution of CX45 puncta fluorescence intensity for 6 SAN subregions and atrial tissue (AT). **F)** Distribution of CX45 puncta area for the same subregions. Outlined boxes highlight CX45 plaques with high fluorescence intensity and large area. All quantifications were pooled from three hearts, one 240 μm x 240 μm image from each heart. (H: head, M: middle, T: tail, A: atrial, central: peri: peripheral, AT: atrial tissue; cent)

Collectively, the CX isoforms defined distinct low electrical coupling domains in the central head and tail subregions and high conductance coupling domains in the mid-central, head-peripheral, and tail-peripheral subregions. Low electrical coupling supports insulation of the centrally located leading pacemaking sites, while enrichment of CX40 in the central middle supports fast conduction between the leading pacemaking sites at the SAN head and tail. Weighing single-channel conductance of CX isoforms, high-conductance CX43 enriched at the head periphery and both CX40 and CX43 enriched at the tail periphery predict strong coupling in those regions, supporting the presence of conduction exit paths in the head and tail peripheries.

## DISCUSSION

The SAN is a comparatively small yet durable current source coupled to large electrical sinks, including both the FBs embedded within it and the ACMs surrounding it, yet it sustains robust pacemaking and pacing functions.^1,2^ This ability has been attributed to an optimal balance between the inter-tissue and intra-tissue current source-sink relationships. Source-sink relationships depend on the intrinsic cell phenotype, cell composition, cell-cell communication, and organization. The PCM phenotype has been shown to differ along the SAN head-to-tail axis.^3,11^ Studies in the mouse SAN have further linked head-to-tail axial differences in cellular energetics, microvascular organization, and stochastic resonance to regional differences in automaticity.^5,12,13^ In this study, we investigated head to tail regional differences in cellular properties and multicellular cluster organizations, including PCM organization and gap junction distribution, that contribute to current source-sink relationships favoring pacemaking and pacing functions.

### SAN Micro-Organization and Its Implications for the Source-Sink Relationship

Previous studies have either analyzed transverse SAN sections or histological staining in longitudinal sections, resulting in limited structural and anatomical information.^4,6–9^ Functionally, optical mapping of *ex vivo* rat and human SAN preparations have shown that leading pacemaking sites are not fixed, but rather switch between two discrete sites localized near the SVC and IVC, with the superior site dominating at fast automaticity frequency >81 bpm and the inferior site at <66 bpm.^3^ The frequency differences have been attributed to regional differences in intrinsic PCM phenotype and cellular organization, which both influence the intra-tissue coupling among PCMs and between PCMs and FBs.^11^ To pace the neighboring atrial tissue, the excitation wavefront exits the SAN via conduction paths, which are located near the SVC, inter-atrial septum, and extending laterally into the atrial myocardium.^4,7,14^ Inter-tissue electrical coupling from the central through peripheral SAN to the atrial myocardium must be balanced to maintain both pacemaking and pacing functions. Insufficient coupling, or excessive electrical insulation of the PCMs from the ACMs, may preserve robust pacemaking in PCMs but prevent pacing of ACMs, whereas excessive coupling may electrotonically overload the PCMs, suppressing their pacemaking and, consequently, pacing function. While studies have uncovered gross anatomical and electrophysiological insights into the SAN, details of the intra- and inter-tissue current source– sink mechanisms linking cell properties, micro-anatomical structure, and cellular organization to proper SAN function remain unclear.

Building on our prior work elucidating the micro-organization of the SAN in the longitudinal orientation,^11^ we quantitatively analyzed the regional differences along the SAN head-to-tail axis. A pseudo-colored image of HCN4 immunostaining showed the superior SAN arching outwards toward the VC with elevated HCN4 expression close to this interface that then gradually transitioned towards the SAN-CT interface. This longitudinal pattern agrees with HCN4 staining in transverse sections used for 3D computational reconstruction of the rabbit SAN.^9^ Consistent with our previous findings,^11^ head and tail PCM clusters are both similarly arranged with lower aspect ratio and less directionality as well as higher HCN4 expression than the mid-region (Figure 6B, 6C). However, the larger cluster size and higher HCN4 expression in the head than the tail region suggest that the head is a stronger current source that can elevate pacemaking strength through both the intrinsic PCM phenotype and multicellular cluster organization, which favor more PCM-to-PCM contact and electrical coupling (Figure 6B, 6C, 6F). These findings provide a structural basis for the localization of the leading pacemaking sites at the head and tail regions in optical mapping studies, with a faster frequency of automaticity at the superior site than the inferior site.^3^ In contrast, large and HCN4-low PCM clusters in the mid-region, in conjunction with high aspect ratio and cluster alignment in the direction of the head-to-tail axis, structurally favors rapid conduction over spontaneous electrical impulse initiation (Figure 6F, 6G). This may provide a high conductance corridor between the head and tail pacemaking sites, enabling quick changes in heart rate via dynamic shifts in the leading pacemaker location.^3^ Synchronization of the two pacemaking poles has recently been revealed to be more complex than entrainment alone. Regionally graded subthreshold voltage fluctuations in the intact node set diastolic coherence through a stochastic resonance mechanism, with each pole operating near its own optimum.^13^ On this view, a high-conductance mid-region corridor would coordinate two independently tuned oscillators rather than simply capturing the slower pole by the faster one.

**Figure 6.**
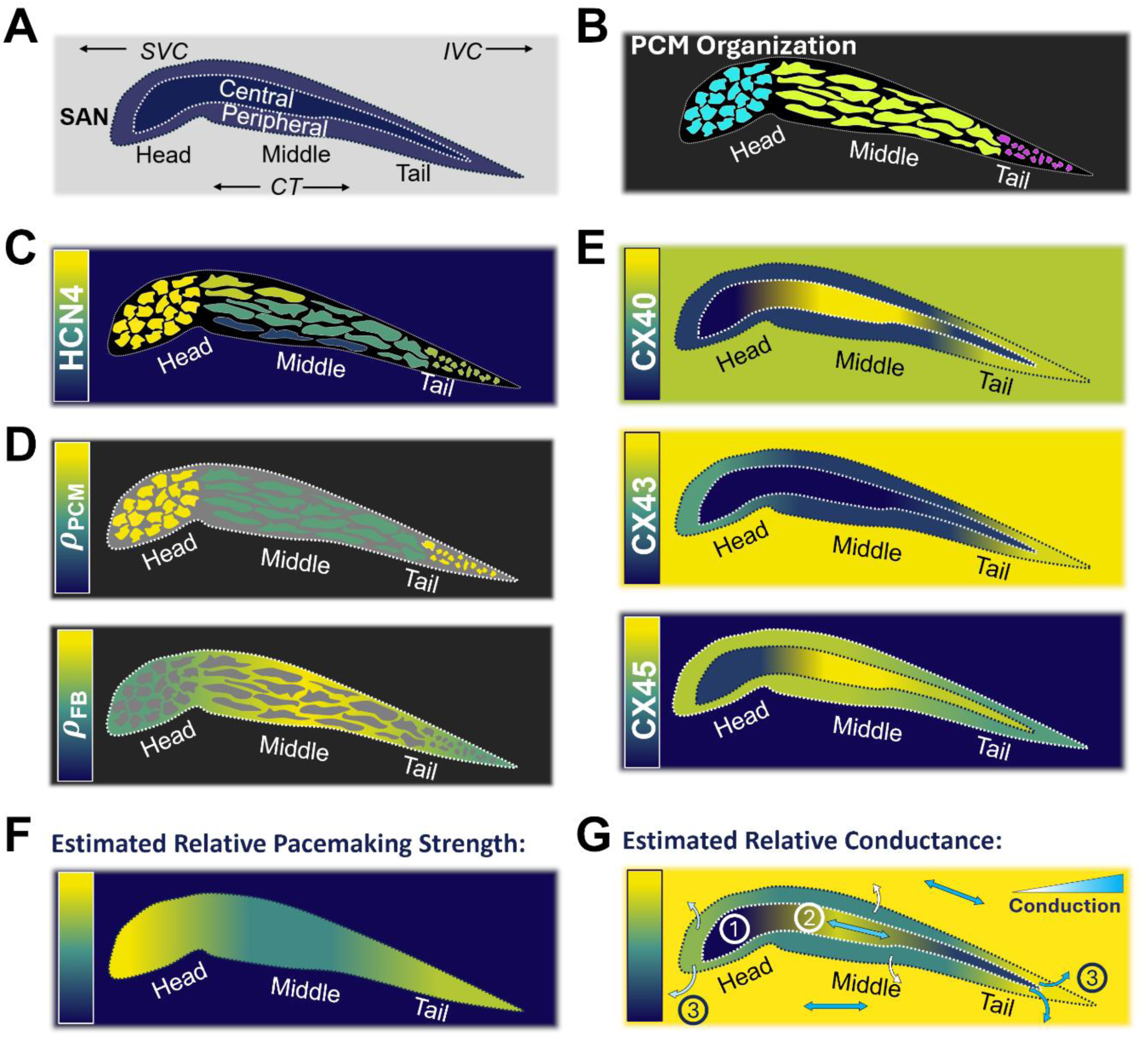
Summary diagrams of the SAN organization. **A)** Diagram of the six subregions— head-central, head-periphery, mid-central, mid-periphery, tail-central and tail-periphery— analyzed in the SAN. **B)** Illustration of the PCM cluster organization and size in the head, middle and tail regions. **C)** Pseudocolor map of relative HCN4 expression across SAN regions. **D)** Pseudocolor maps of relative local PCM density (*ρ_PCM_*) and local FB density (*ρ_FB_*) in the SAN regions. **E)** Pseudocolored maps of relative CX expression for each isoform across SAN subregions. **F)** Pseudocolor diagram of estimated relative pacemaking strength across the SAN. **G)** Pseudocolor diagram of estimated relative conductance magnitude, with arrows indicating the direction of conduction inferred from PCM cluster alignment and aspect ratio. Key insights highlighted include: 1) low conductance in the central head subregion, 2) high conductance in the central middle subregion, and 3) high conductance radially away from the SAN at the head and tail peripheries.

### Cellular Analyses of the SAN Head, Middle, and Tail

Moving down the spatial scale from cluster analysis, we examined cellular compositions within clusters of the head, middle, and tail regions. PCMs in the SAN function as the current source, whereas FBs electrically coupled to PCMs through gap junctions can act as a current insulator, an electrotonic sink, or an electrotonic source, depending on the strength of coupling and the phase of the action potential.^2^ With minimal PCM-FB electrical coupling, FBs are effectively an electrical insulator. With adequate PCM-FB electrical coupling, even though FBs can supply electrotonic depolarizing current to PCMs in phase-4 of action potentials that mildly facilitates PCM depolarization towards action potential activation threshold, FBs generally behave as a current sink that drains PCMs electrotonically once action potential upstroke surpasses the resting membrane potential of FBs.^15–19^ Considering that a large current sink relative to the current source can effectively reduce the frequency of automaticity or completely silence automaticity in PCMs, statistically higher local PCM density, in conjunction with a trend toward lower local FB density, equate to a strong current source and weak current sink in the SAN head and tail, providing a compositional condition that supports the leading pacemaking sites located at the SAN head and tail (Figures 6D, 6F). While comparable in local cell densities, the weaker pacemaker profile with lower HCN4 expression and smaller PCM clusters in the SAN tail than the head is consistent with slower pacemaking frequency in the inferior leading pacemaking site as reported in optical mapping studies.^3,20^ A combination of strong pacemaking cellular phenotype and cluster arrangement, including high HCN4-expressing PCMs, high PCM density, low FB density, and large PCM cluster organization forming an intra-tissue source-sink (PCM-FB) relationship that maximizes pacemaking function and minimizes electrotonic load, are critical for the emergence of a dominant, fast leading pacemaking site at the SAN head (Figures 6A-6D, 6F). This structural asymmetry has parallels in the blood supply and energetics of the node. In mouse, microvascular density is higher in the superior than in the inferior node. PCMs in the SAN head benefit from close proximity to vessels, with beat-locked ATP production at a higher gain and consequently faster intrinsic automaticity.^5^ In contrast, PCMs in the SAN tail fire at rates constrained by operating nearer an energy deficit.^12^ Whether the head-to-tail gradients in PCM density and cluster size reported here align with comparable vascular and metabolic gradients in the pig remains to be tested.

### Numbers of PCMs and FBs within the SAN

To date, the literature on the SAN, hiPSC-derived PCMs, and engineered biopacemakers frequently cites ∼5,000 PCMs residing in the SAN, reported by Bleeker *et al*., in 1980; however, that estimation of a rabbit SAN was not derived from cell quantification.^21^ A similar case is the widely cited 100 billion neurons and 1 trillion glial cells in the human brain that became an accepted “fact” through repeated citation but was later shown to be substantially lower by direct quantification.^22^ To determine the number of PCMs and FBs within a rabbit SAN of ∼1 mm^3^ in volume,^23,24^ we estimated the total numbers of PCMs and FBs residing in the SAN using regional cell densities and assuming that the head and tail regions each occupy ∼25% of the total SAN space, that the thickness corresponds to an optical section of 11 μm (based on the numerical aperture of the objective and the index of refraction of the mounting medium), and that cross-species differences in cell density are minimal. The SAN was quantitatively estimated to consist of ∼45,000 PCMs and ∼73,000 FBs, exceeding reported PCMs in the rabbit SAN by 9-fold. Extending this cell estimation to human SANs, if we assume for simplicity a cylindrical SAN volume that is 5 mm in diameter and 20 mm in length, the calculated cell composition in the SAN based on the regional densities equates to ∼1.9 x 10^7^ PCMs and 2.9 x 10^7^ FBs. These more accurate estimates of PCM and FB numbers and densities are valuable insights for engineering biopacemakers and for interpreting electrophysiological studies.

### Functional Implications of the Expression and Distribution of the Different Isoforms of CX within the SAN

The strength of electrical coupling between PCMs to PCMs, ACMs, and FBs is mediated by CX-based gap junctions that serve as another current source-sink modulator. Three major CX isoforms have been reported in the heart, with CX40 predominantly in the atria, CX43 in the ventricles, and CX45 in the SAN.^25,26^ Analyses of CX40, CX43, and CX45 distributions in porcine SANs divided into 6 subregions are summarized in pseudocolored effective conductance maps that were calculated and normalized based on the CX puncta density, puncta size, and previously reported single-channel conductance for each isoform—CX40 at ∼220 pS, CX43 at ∼120 pS, and CX45 at ∼40 pS^27,28^—with the assumption that every punctum formed functional gap junctions (Figures 6E, 6G). Effective conductance was estimated as follows:

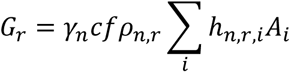

where *G_r_* is the estimated effective gap junction conductance in region *r* (per 1000 µm²), *γ_n_* is the single-channel conductance of connexin isoform *n*, *c* is the channel packing density, *f* is the fraction of open channels (assumed 1), *ρ_n_*_,*r*_ is the mean puncta density of connexin isoform *n* in region *r* (puncta per 1000 µm²), ℎ is histogram distribution of puncta size in bin *i*, and *A_i_* is the punctum area. The effective conductances for each CX isoform were then normalized to facilitate heatmaps generation. Our data are in general agreement with reported CX isoform distribution, including high CX45 and low CX43 in the SAN; however, we found that the CX isoform distribution is highly heterogeneous in the porcine SAN, differing not only in the head-to-tail axis but also radially from central to periphery.

Low expression of high-unitary-conductance CX40 and CX43 in PCMs residing in the central SAN head and tail indicates weak electrical coupling, mediated predominantly by low-unitary-conductance CX45 in these leading pacemaking regions. This coupling profile may protect the current source from electrotonically overloading by the surrounding current sink. In contrast, CX40 is remarkably abundant in PCMs present in the central-middle SAN similar to that in the atrial region, with high CX40 puncta density and large plaque size. The abundance of CX40 suggests a principal conductance highway within the central middle subregion. Notably, lower CX40 and CX43 expression in FBs within this subregion may limit their current-sink suppression on conduction, further supporting a fast-conducting connection between the two leading pacemaking sites. The magnitude and direction of conductance in the SAN central mid-subregion is further enhanced by large CX40 puncta plaque at the intercalated discs and the head-to-tail axial alignment with high aspect ratio PCMs. The CX expression and cell organization infers intra-tissue anisotropic conduction across the SAN, especially in the mid-region, that is consistent with reported elliptical isochronal activation maps of SANs acquired via optical mapping.^3^

CX40 and CX43, both high-unitary-conductance isoforms, are abundant in PCMs of the SAN tail peripheral subregion, indicating a highly conductive region. Although lower in conductance than the tail periphery, the SAN head periphery is presumably more conductive than the central subregion at the two ends due to higher CX43 and CX45 expressions. This inter-tissue coupling pattern reveals a radial conductance gradient—from low central to high peripheral conductance— in the SAN head and tail. This gradient points to an electrically insulated core surrounded by a well-coupled periphery, which may create a low-resistance pathway that preferentially directs current outward from the SAN and supports its ability to pace the atrial tissue. These peripheral domains therefore provide a plausible structural basis for SAN conduction exit pathways detected along the SAN-atrial interface, facilitating excitation wavefront propagation out of the SAN, while preventing the atrial tissue from electrotonically suppressing the pacemaking function.^7,14,29^ In the tail, lower HCN4 and greater peripheral CX40 and CX43 expression predict lower current source and stronger coupling, consistent with lower intrinsic pacemaking strength than the head but supporting effective conduction at the peripheral exit. The relative enrichment of CX43 and CX45 in the central tail is also qualitatively consistent with reported faster inferior than superior intranodal conduction (8.7 vs. 6.5 cm/s).^4^

Conduction insights are summarized by the estimated scalar total effective conductance value for each subregion based on the integrated CX isoform conductance with preferential conducting direction indicated by the arrows that were inferred from PCM cluster alignment and aspect ratio (Figure 6G). Of note, the pseudocolor conductance maps did not include correction for a nonlinear increase in channel access resistance with increase in puncta size. However, with correction based on numerically simulated puncta packing density to the access resistance relationship,^30^ the estimated conductance trend in the SAN remains the same (pseudocolor conductance map not shown). Additionally, the conductance pseudocolor maps were not corrected for critical plaque size of 200-400 channels for channel opening.^31^ Given a gap junction channel size of 100 nm^2^, the smallest puncta of 0.2 μm^2^ in our images is well above the critical plaque size threshold of 0.04 μm^2^. Other notable connexin patterns observed in our images include the co-localization of only CX40 with HCN4 at the intercalated discs between PCMs in the central mid-region, whereas in the surrounding ACMs that express low HCN4, both CX40 and CX43 co-localized with HCN4 at the intercalated discs. We also did not observe large (≥ 5 µm²) CX45 plaques in any cell type. Although all three CX isoforms were detected in SAN FBs, detailed analysis of puncta size and electrical coupling between FBs and PCMs, which may modulate the intra-tissue electrical coupling, remain unresolved. Formation of heterotypic channels or heteromeric connexons from combined CX isoforms was also not investigated in our current study due to the limitation of antibody combination. Gap junctions provide not only electrical coupling but also direct chemical coupling for any molecules < 1 kDa.^30^ Heterotypic channels have been shown to serve as voltage-sensitive intercellular valves, such that small static trans-junctional voltages or high frequency fluctuations can engage the regulatory valve, resulting in unidirectional signaling and/or electrotonic conduction.^32^ Therefore, the effects of heterotypic channels involved in homocellular and heterocellular coupling on SAN pacemaking and pacing functions warrant further investigation.

## CONCLUSIONS

Collectively, our data provide new structural and organizational insights, revealing that the SAN has a heterogeneous, region-specific coupling architecture supporting pacemaking and pacing functions. Intra-tissue coupling is predicted to 1) electrically insulate the central SAN head and tail, thereby protecting their function as the leading pacemaking sites, and 2) provide an electrically well-coupled central mid-region that serves as a high-conduction nexus coordinating the superior and inferior leading pacemaking sites for fast heart rate changes. In contrast, inter-tissue coupling is predicted to support efficient conduction through the exit pathways at the SAN head and tail peripheries, allowing these peripheral subregions to couple with the atrial myocardium and support proper SAN pacing function. This spatial specialization in cellular phenotype, organization, and electrical coupling expands our understanding of the regional SAN pacemaking and pacing functions along the head-to-tail axis. These findings represent an important step toward linking SAN structure to function and help inform the development of biopacemakers, both as experimental models and regenerative therapies, as well as strategies for pharmacological therapies.

## ACKNOWLEDGEMENTS

Hillary K. J. Kao: experimental design, data collection, data analysis, data interpretation, and manuscript preparation

Regan Smithers: data collection, data analysis, data interpretation, and manuscript preparation

David H. Ramos-Rodriguez: data interpretation

Kaitlin Van Brusselen: data collection

Rableen K. Kaleka: data collection

David Cameron: data collection

Jan A. Nolta: data interpretation

Kyle Fink: resources

J. Kent Leach: resources

L. Fernando Santana: data interpretation

Nipavan Chiamvimonvat: resources and data interpretation

Deborah K. Lieu: experimental design, data analysis, data interpretation, manuscript preparation, supervision, resources, and funding acquisition

## NONSTANDARD ABBREVIATIONS AND ACRONYMNS

ACM: atrial cardiomyocyte
CT: crista terminalis
CX: connexin
EC: endothelial cell
FB: fibroblast
HCN4: hyperpolarization-activated cyclic nucleotide-gated 4 channel
PCM: pacemaking cardiomyocyte
SAN: sinoatrial node
TNT: troponin T
VC: vena cava
VIM: vimentin

## SUPPLEMENTAL FIGURES WITH FIGURE LEGENDS

**Figure S1.**
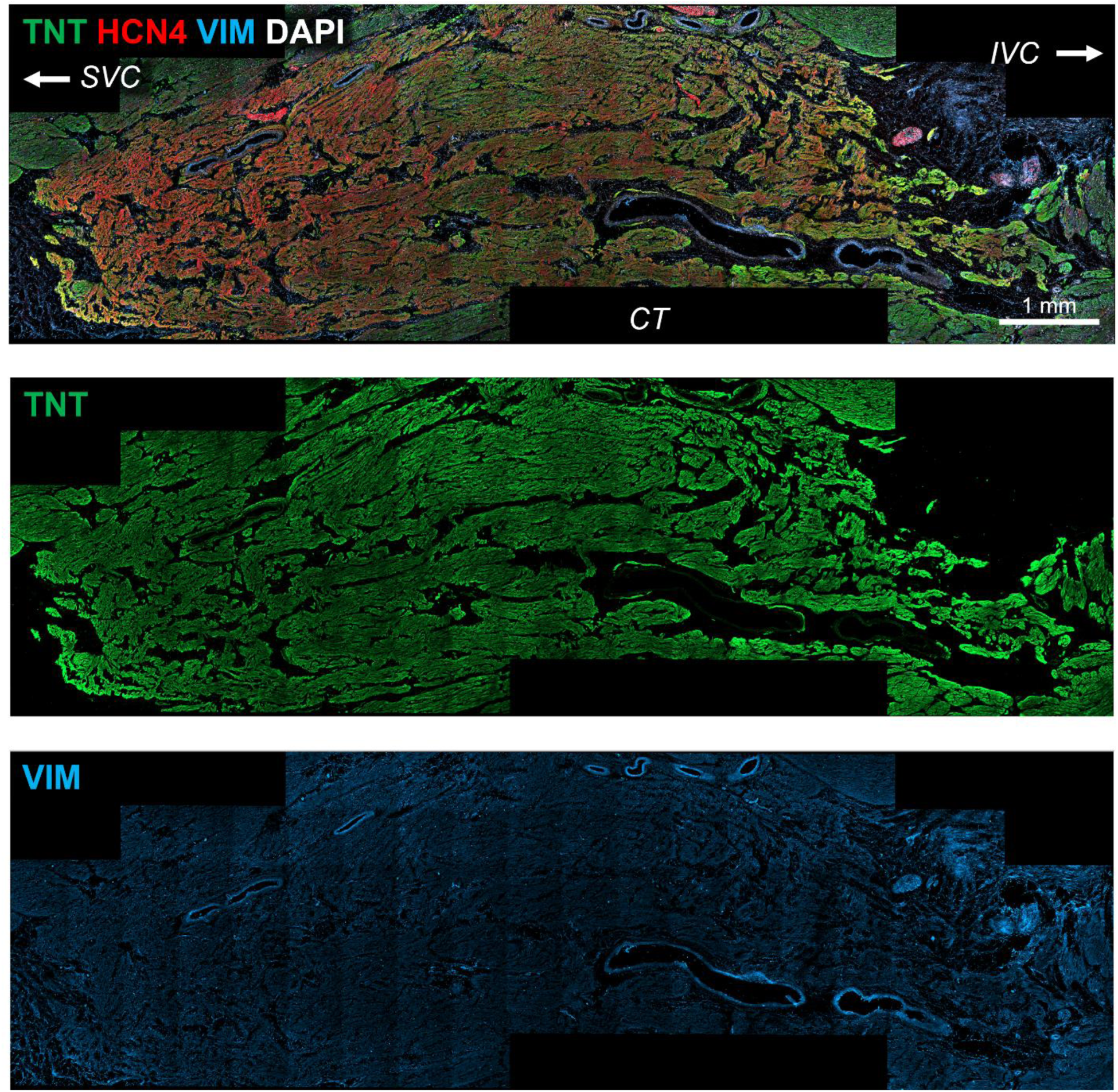
Representative SAN section immunostained for TnT, HCN4, VIM, and DAPI. These are the original fluorescence images presented in Figure 1.

**Figure S2.**
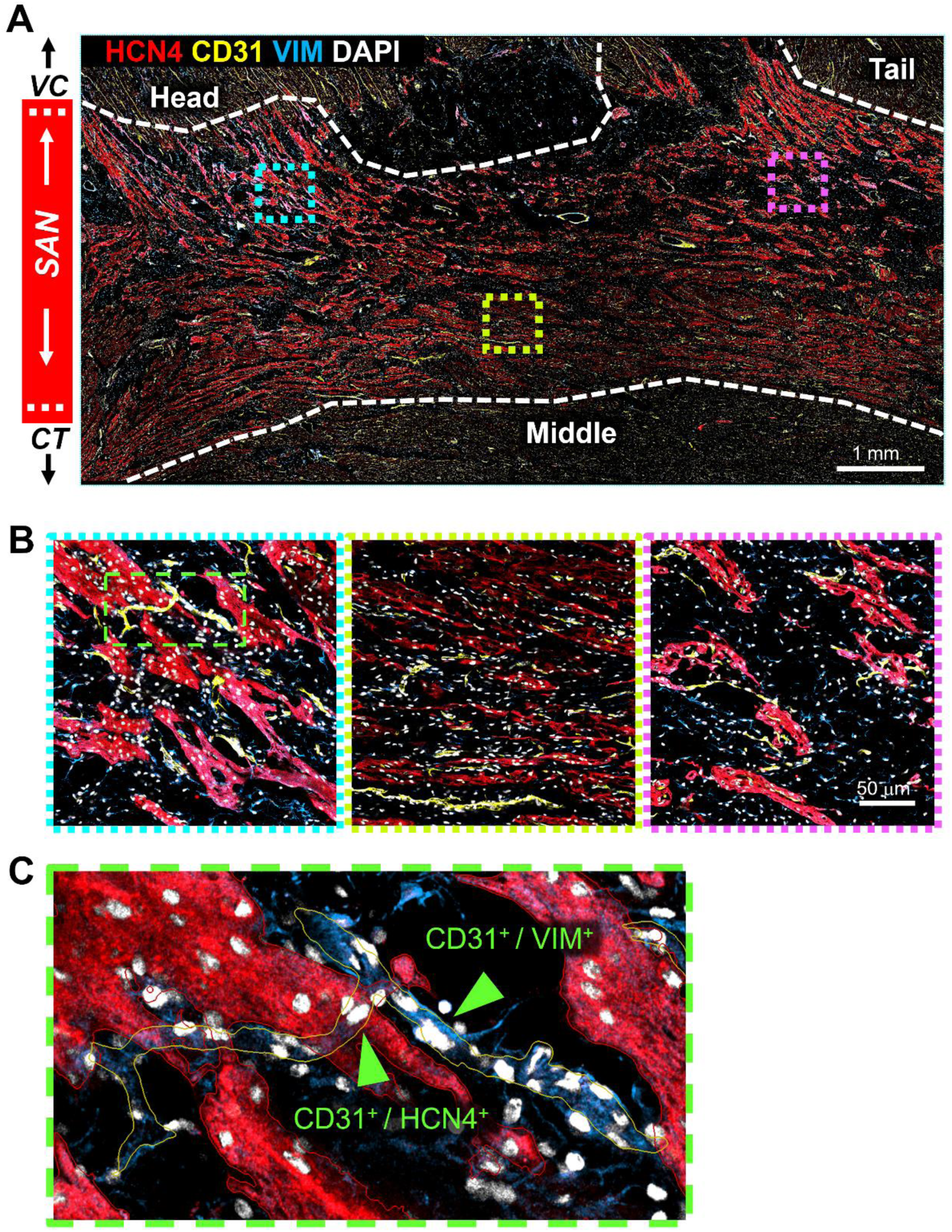
Endothelial cells (ECs) in the porcine SAN. **A)** SAN cryosection immunostained with HCN4, CD31, and VIM to identify, respectively, PCMs, ECs, and FBs. Dashed white lines indicate the SAN-atrial borders. **B)** Magnified view for region in the outlined boxes in the head, middle, and tail regions showing in panel A. **C)** Zoomed in image for the green outlined region in panel B showing CD31^+^ endothelial nuclei (green arrowheads) in the HCN4^+^ or VIM^+^ structure that were misclassified as PCMs or FBs.

**Figure S3.**
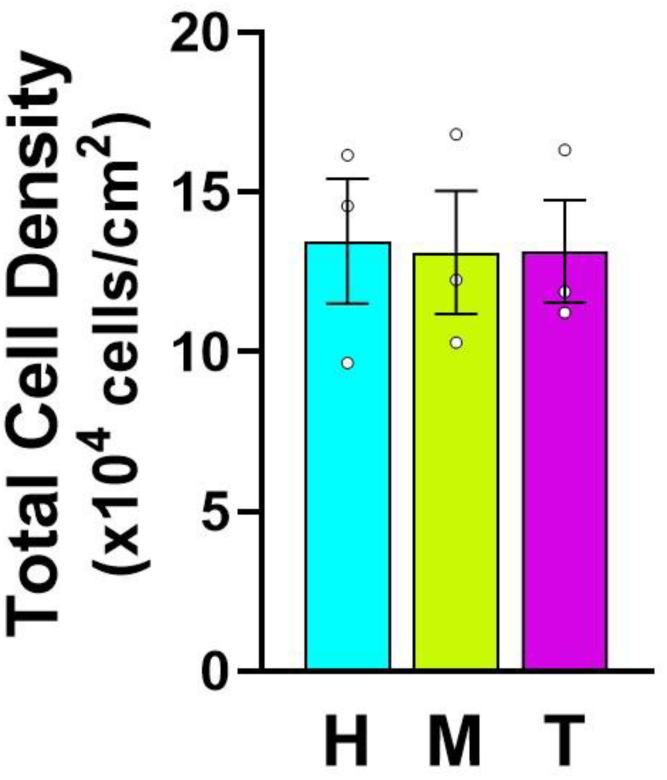
Total cell density in the head (H), middle (M), and tail (T) regions of porcine SANs. Cell density in the regions is shown regardless of cell type.

**Figure S4.**
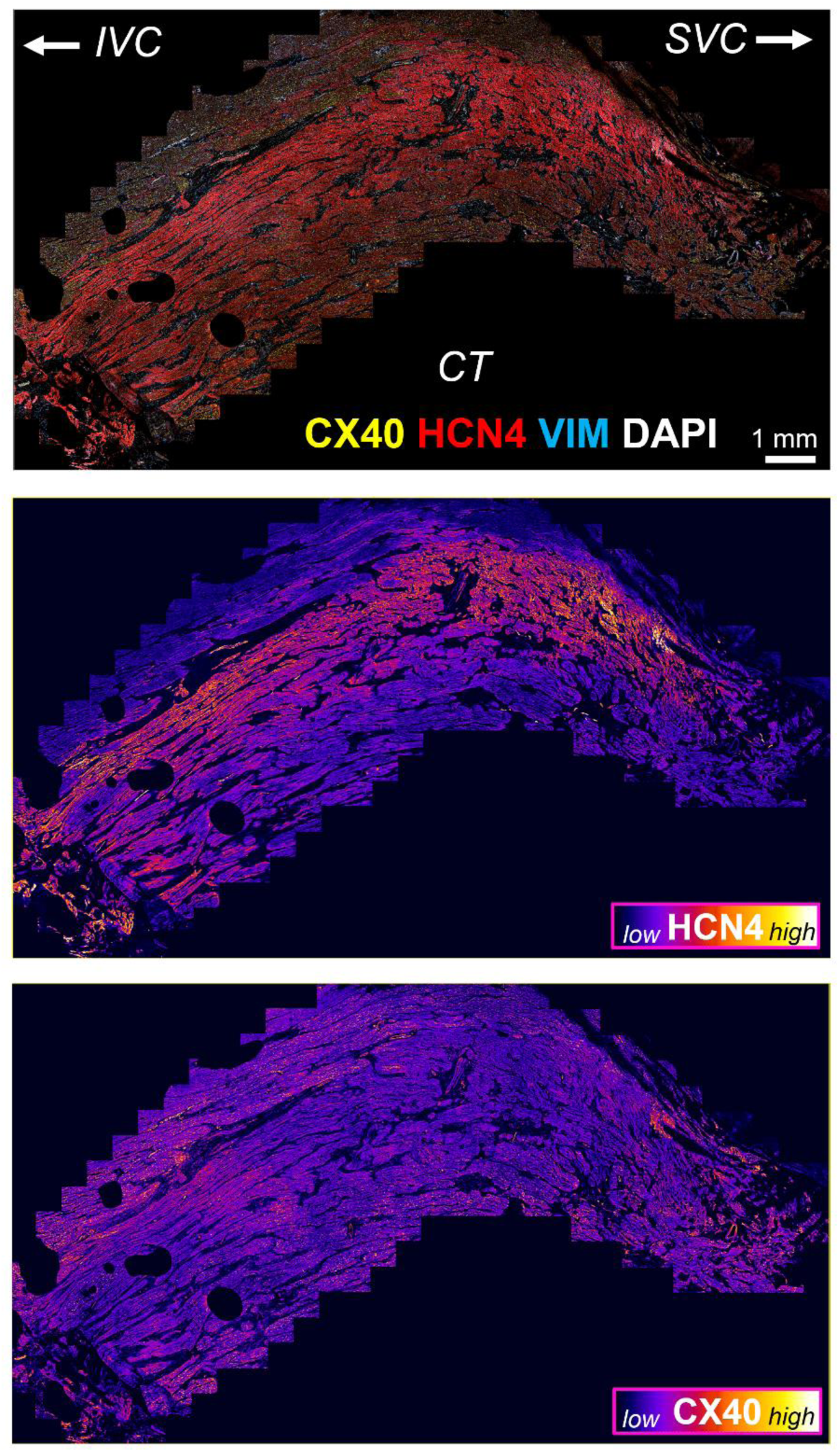
CX40 distribution in the porcine SAN along the head-to-tail axis. Fluorescence image of the SAN immunostained for HCN4, CX40, VIM, and DAPI, with corresponding pseudocolor images showing HCN4 and CX40 expression individually.

**Figure S5.**
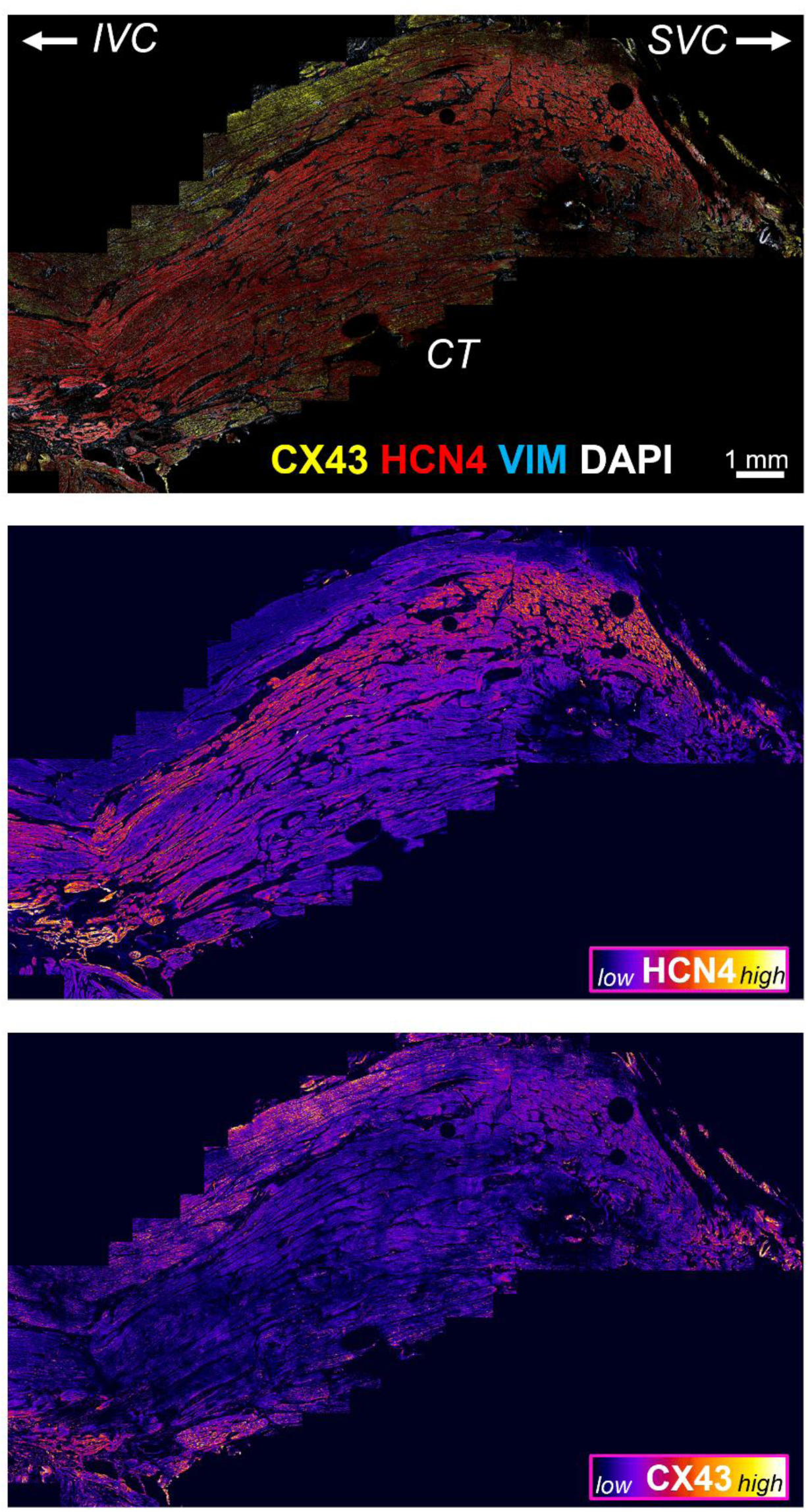
CX43 distribution in the porcine SAN along the head-to-tail axis. Fluorescence image of the SAN immunostained for HCN4, CX43, VIM, and DAPI, with corresponding pseudocolor images showing HCN4 and CX43 expression individually.

**Figure S6.**
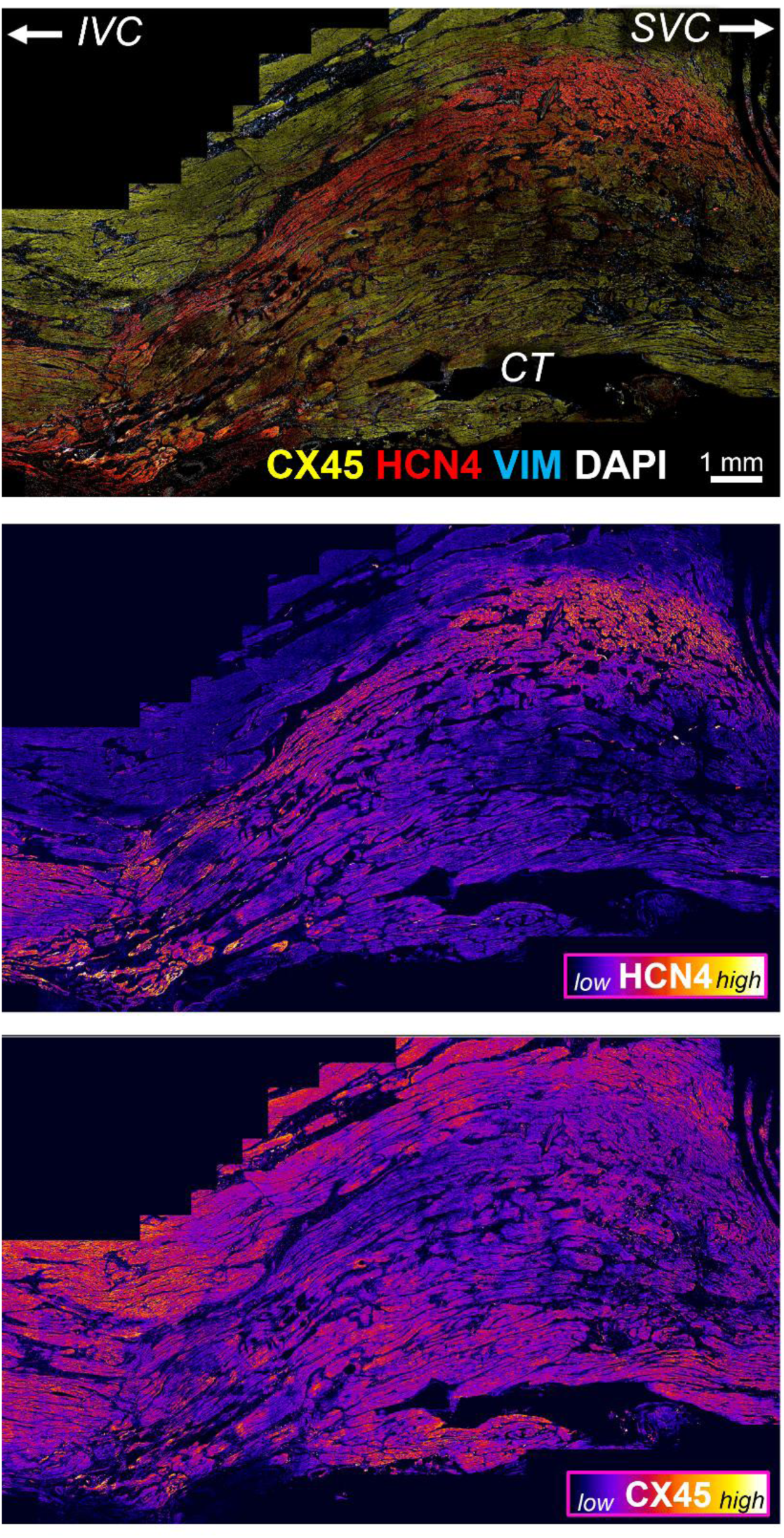
CX45 distribution in the porcine SAN along the head-to-tail axis. Fluorescence image of the SAN immunostained for HCN4, CX45, VIM, and DAPI, with corresponding pseudocolor images showing HCN4 and CX45 expression individually.

**Figure S7.**
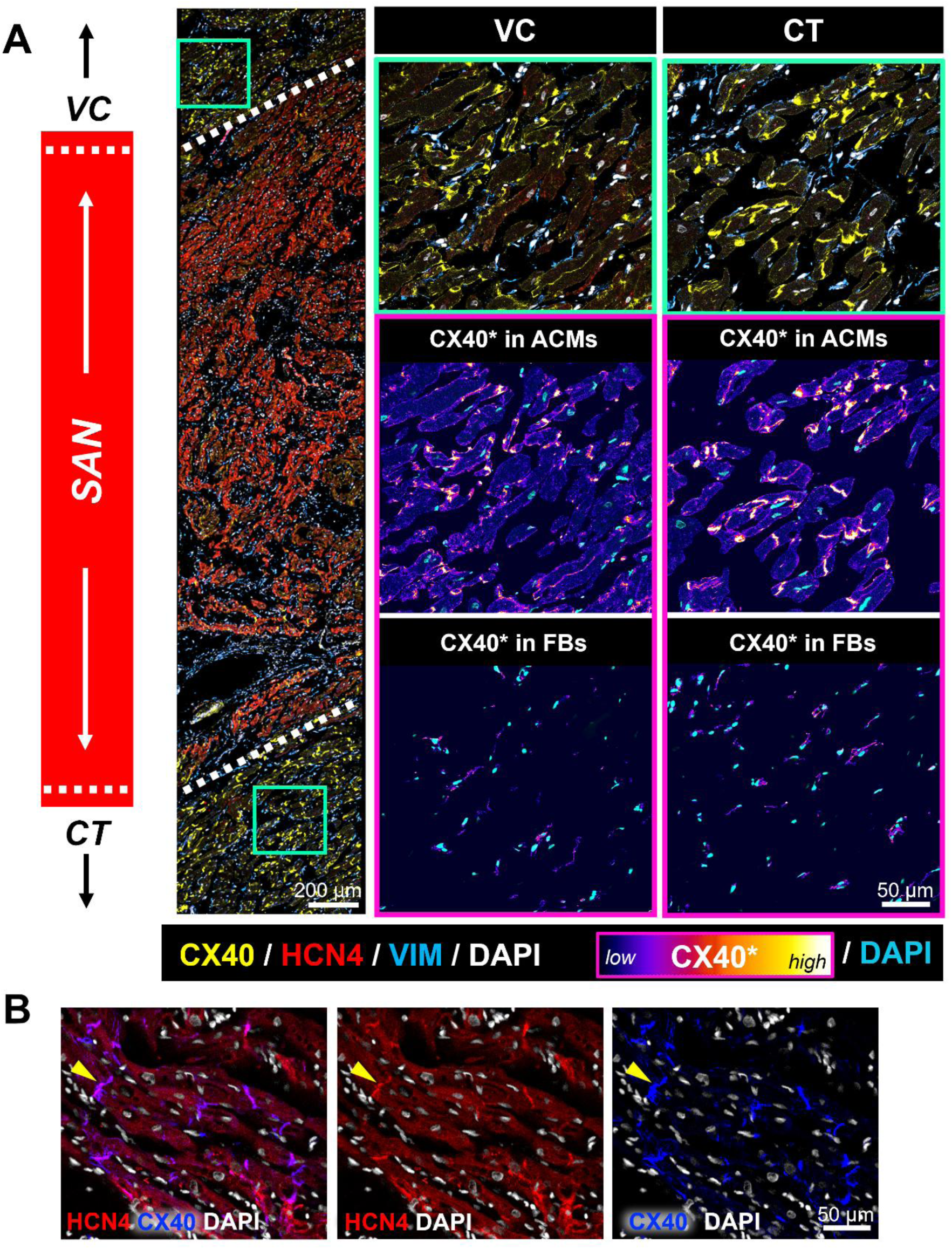
CX40 expression in CT and VC atrial regions. **A)** Representative immunofluorescence images for CX40, HCN4, VIM, and DAPI. Fluorescence image insets show magnified VC and CT atrial regions with corresponding pseudocolor images of CX40 expression in ACMs and FBs. **B)** Magnified fluorescence images of atrial tissue showing co-localization of HCN4 with CX40, as indicated by the yellow arrowhead.

**Figure S8.**
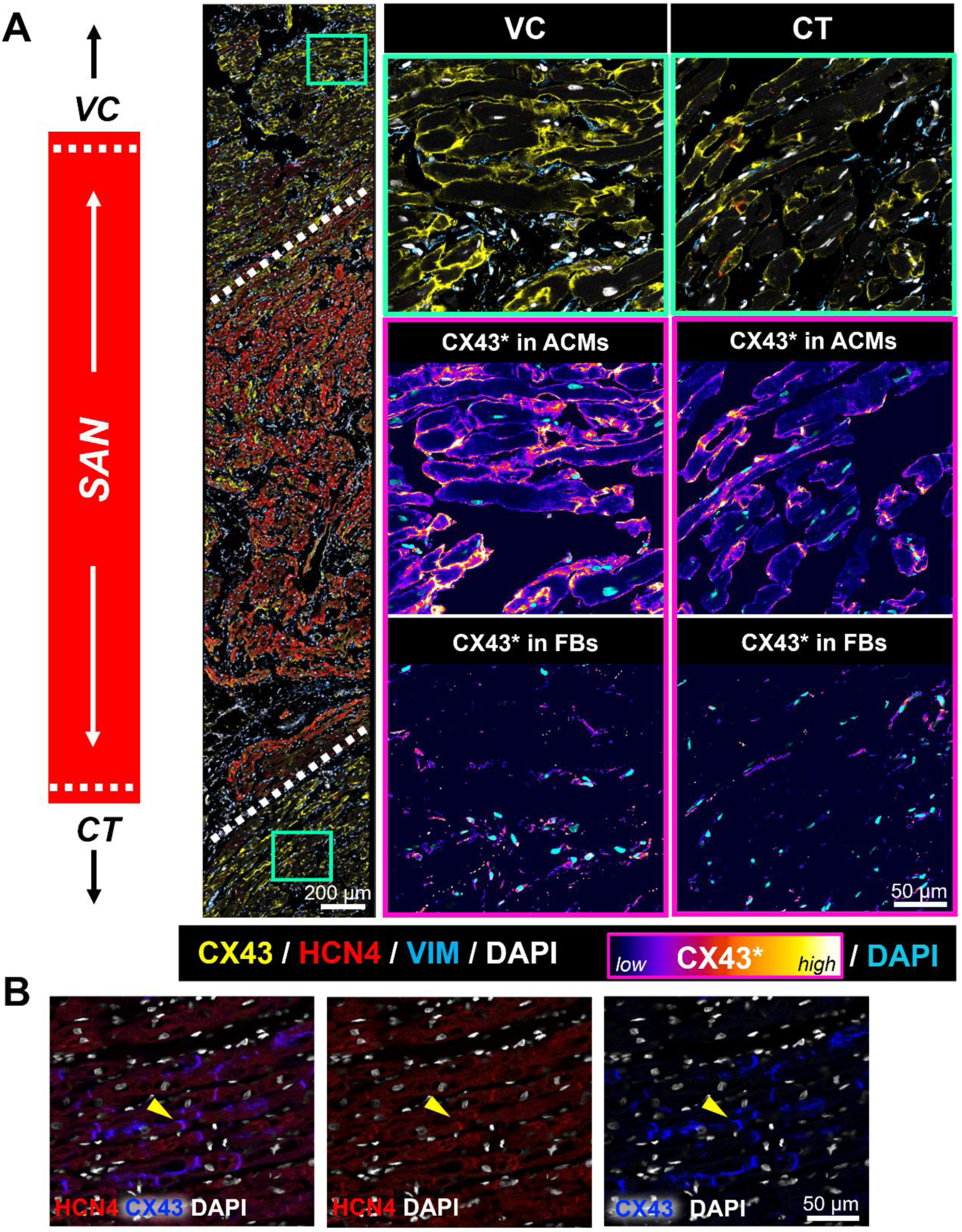
CX43 expression in CT and VC atrial regions. **A)** Representative immunofluorescence images for CX43, HCN4, VIM, and DAPI. Fluorescence image insets show magnified VC and CT atrial regions with corresponding pseudocolor images of CX43 expression in ACMs and FBs. **B)** Magnified fluorescence images of atrial tissue showing co-localization of HCN4 with CX43, as indicated by the yellow arrowhead.

**Figure S9.**
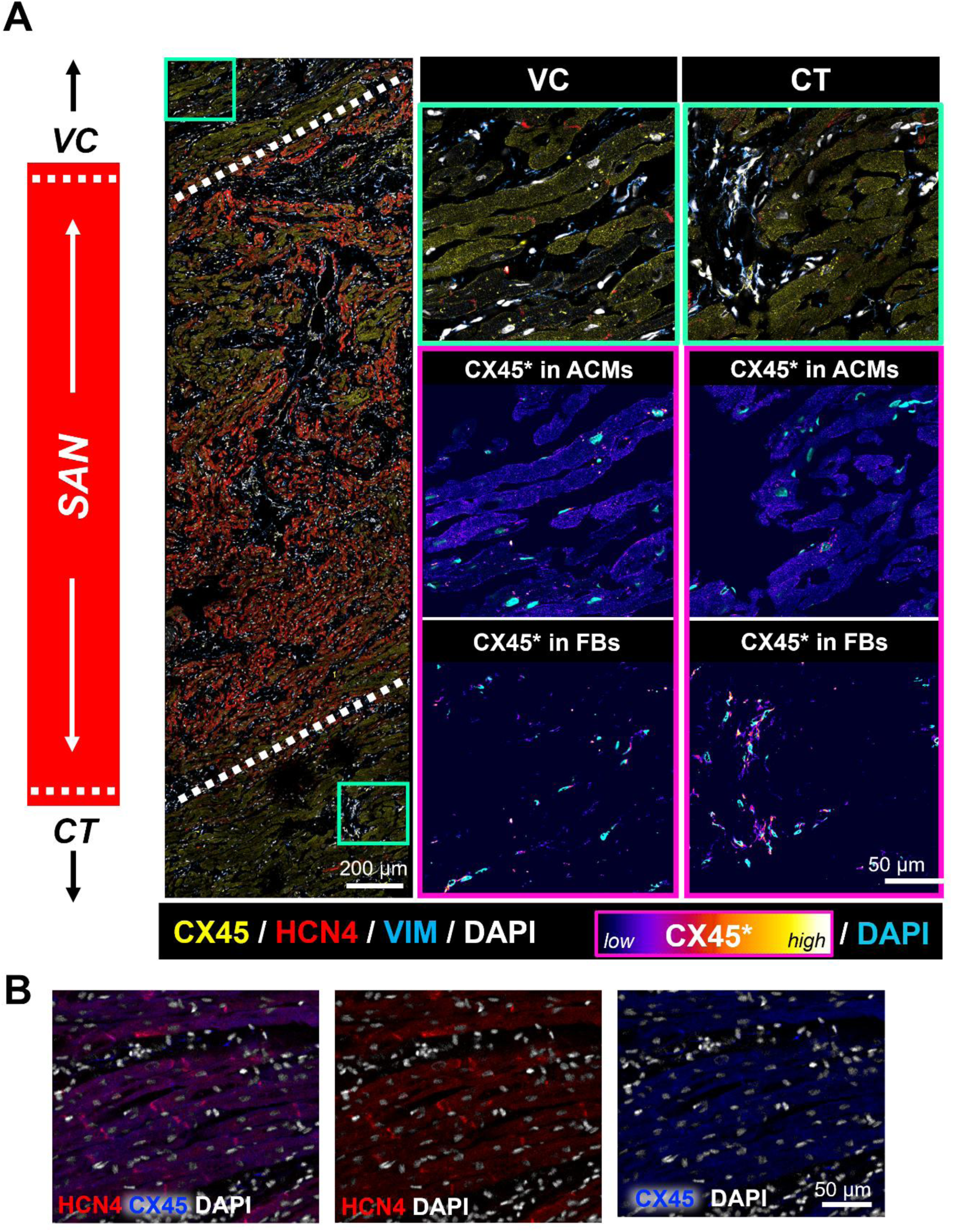
CX45 expression in CT and VC atrial regions. **A)** Representative immunofluorescence images for CX45, HCN4, VIM, and DAPI. Fluorescence image insets show magnified VC and CT atrial regions with corresponding pseudocolor images of CX45 expression in ACMs and FBs. **B)** Magnified fluorescence images of atrial tissue showing absence of co-localization between HCN4 and CX45.

**Table S1.** List of antibodies used in SAN immunostaining.

|  | **Reagent** | **Company Source** | **Catalog No.** | **Dilution Factor** |
|---|---|---|---|---|
| **Primary antibodies** | HCN4 (mouse monoclonal IgG1) | Antibodies Inc. | N114/10 | 1:200 |
|  | cTNT (mouse monoclonal IgG2a) | Developmental Studies Hybridoma Bank | CT3 | 1:200 |
|  | CX40 (rabbit polyclonal IgG) | Invitrogen | 36-4900 | 1:200 |
|  | CX43 (rabbit polyclonal IgG) | Proteintech | 26980-1-AP | 1:200 |
|  | CX45 (rabbit polyclonal IgG) | BiCell Scientific | 00407 | 1:200 |
|  | CD31 (rabbit polyclonal IgG) | Bethyl Laboratories | IHC-00055 | 1:200 |
|  | VIM (chick polyclonal IgG) | Invitrogen | PA1-10003 | 1:5000 |
| **Secondary antibodies** | Alexa Fluor 488 goat anti-mouse IgG1 | Invitrogen | A-21121 | 1:1000 |
|  | Alexa Fluor 546 goat anti-mouse IgG1 | Invitrogen | A-11010 | 1:1000 |
|  | Alexa Fluor 488 goat anti-mouse IgG2a | Invitrogen | A-21131 | 1:1000 |
|  | Alexa Fluor 546 goat anti-rabbit IgG | Invitrogen | A-11035 | 1:1000 |
|  | Alexa Fluor 647 goat anti-chick IgG | Invitrogen | A-21449 | 1:1000 |

